# Mixed signals: heat stress alters chemical profiles and increases same-sex sexual behaviour in an insect

**DOI:** 10.64898/2026.09.21.753124

**Authors:** Solène A. Morelle, Johannes Stökl, Nathan W. Bailey, Sandra Steiger, Natalie Pilakouta

## Abstract

Rising temperatures linked to global climate change can disrupt the chemical communication that animals rely on for reproduction. Arthropods are particularly vulnerable due to their limited ability for thermoregulation and their reliance on cuticular hydrocarbons (CHCs), which have a dual role in social communication and heat protection. Heat-induced changes in cuticular composition may therefore lead to mate recognition errors. We tested this in the burying beetle *Nicrophorus vespilloides*, by exposing males to either benign temperatures (20°C) or a simulated 3-day heatwave (26°C). We recorded the frequency of same-sex sexual behaviour under these contrasting thermal conditions and examined associated changes in the males’ cuticular profiles. Our results show a higher rate of same-sex mounting under heat stress, with the CHCs of heat-stressed males shifting toward more saturated, longer-chained compounds and more female-like compounds. We also found a positive association between chain length and the frequency of same-sex mounting, suggesting a trade-off between the social communication and waterproofing functions of CHCs. Overall, our findings provide evidence for a causal effect of heat on same-sex sexual behaviour, which may be partly explained by changes in cuticular lipids. Such disruption of chemical communication could have major consequences for animal reproduction in a warming world.

## Introduction

In light of global climate change, there is increasing concern about the effects of rising temperatures on animal reproduction[1,2]. Heat stress can reduce fertility and fertilisation success[3,4], lower fecundity, hatching success, and offspring fitness[5–7], and impair reproductive behaviours such as mating latency, discrimination and choosiness, and parental care[8–10]. Despite a growing body of research on this topic, a crucial component of reproduction that requires further research is the effect of warming on chemical communication[11].

Animals employ a wide range of communication modes during reproduction, but chemical communication is arguably the most common mode, being shared across all lifeforms, from single-celled organisms to the most complex animals[12]. Every aspect of chemical communication, from the production of an information conveying chemical, or infochemical[13], by a sender to the behavioural response of a receiver, is susceptible to elevated temperatures[14]. These effects are general to chemical communication but also apply to the pheromonal signalling used in reproduction. Heat stress can influence the enzymatic activities underlying infochemical biosynthesis, thereby altering the ratio of compounds within a chemical blend[15], including pheromone mixtures[16]. Elevated environmental temperatures can also alter the timing and quantity of volatile infochemical release[17] and shift blend composition as compounds evaporate at different rates depending on molecular weight[18–20]. Because behavioural responses often depend on the specific combination of compounds, these effects may reduce the efficiency of communication, which can be particularly detrimental in reproductive contexts[11,21,22]. On the receiver end, a higher temperature can influence the chemosensory receptors, neuronal signalling, and brain processing through temperature-dependent biochemical pathways[23–25]. As a consequence of these disturbances, warming can leave organisms more or less responsive to infochemicals, including pheromones, thereby impairing the coordination of reproduction[26,27].

These effects may be especially pronounced in arthropods, which rely heavily on chemical signalling to mediate reproduction[28,29] and, as generally small-bodied ectotherms, are especially sensitive to thermal stress[30,31]. Indeed, the arthropod cuticle is covered by a layer of lipids composed mainly of cuticular hydrocarbons (CHCs) such as alkanes, alkenes and methyl-branched alkanes, but also including oxygenated compounds such as aldehydes[32]. These CHCs play a key role in social communication, ranging from nestmate and colony recognition, to sex pheromones and antiaphrodisiacs[33–35]. Importantly, cuticular profiles are not static but are shaped by diet, physiological condition, and environmental factors, especially temperature[36,37], with thermally induced changes emerging as rapid plastic responses within as little as a day[38]. Understanding how thermal variation alters the arthropod cuticle is therefore critical for predicting behavioural responses to warming and forecasting population vulnerability.

As elevated temperatures increase the risk of cuticular water loss, they lead to a shift in the cuticular profile to improve waterproofing[39]. This creates a trade-off between desiccation resistance and mate signalling: higher temperatures favour greater investment in long-chain, saturated CHCs, which form a viscous, waterproofing layer[40], whereas effective signalling usually depends on compounds fluid enough to be perceived[41,42]. Indeed, the two CHC classes most consistently implicated in sex recognition are alkenes and methyl-branched alkanes[43–45], whose double bonds and branch points lower their melting temperature, whereas the waterproofing function is associated with saturated straight-chain n-alkanes[46]. Because a functional CHC layer must provide both waterproofing and signalling simultaneously, warming can shift its composition in ways that alter the information available to receivers, potentially disrupting species and/or sex recognition[41]. Heat-induced changes in cuticle composition could therefore disrupt arthropod reproduction at the very first step of mate discrimination. Such disruption may have several behavioural consequences: individuals may fail to identify an appropriate mating partner[46,47], become less able to assess potential mate quality[40], or even lose the ability to reliably discriminate between the sexes[48]. In the latter case, reduced reliability of sex-specific chemical cues could lead to mate recognition errors, including sexual behaviour directed towards same-sex individuals.

Same-sex sexual behaviour is widespread across animal taxa and may arise from adaptive, neutral, or mechanistic causes[49–51]. One potential mechanistic cause is the breakdown of sex-specific chemical recognition[52]. For example, in *Drosophila*, changes in the chemical information carried by cuticular lipids can lead to same-sex courtship. Genetically ablating the oenocytes that synthesise CHCs, or targeting specific cuticular compounds responsible for sexual dimorphism, produced males that became highly attractive to other males, eliciting same-sex courtship behaviours[53,54]. Consistent with the idea that a change in CHCs can redirect courtship, experimentally applying male-specific compounds to females significantly delays mating, whereas re-adding female-specific compounds attenuates these effects[55]. Such findings indicate that a male exhibiting an altered cuticular profile can become a target of same-sex courtship. This raises the compelling hypothesis that elevated temperatures alter cuticular chemical profiles in ways that reduce the reliability of sex-specific cues, leading to mate recognition errors likely to increase the frequency of same-sex sexual behaviour.

Here, we test whether heat stress leads to shifts in the cuticular profile and alters the frequency of same-sex sexual behaviour. We used the subsocial burying beetle *Nicrophorus vespilloides*, which provides a tractable model system for linking climate change, chemical signalling, and same-sex sexual behaviour. Burying beetles rely heavily on chemical cues for mate recognition and social interactions on the ephemeral vertebrate carcasses on which they breed, where intense competition favours rapid and accurate discrimination of conspecifics and prospective mates[56]. These cues allow them to identify species[57], sex[58], breeding status[59], and previous mating partners[60]. Sex recognition operates at close range: a male assesses an encountered beetle by antennal contact of the cuticle. Because male and female profiles overlap, this discrimination is thought to be imperfect and lead males to mount other males[61], although there is also some evidence that this behaviour may be adaptive (Parlamis et al., in prep). Another advantage of this model system is that reproduction and parental care are known to be highly sensitive to thermal stress (e.g.[62–64]), making this an ideal system in which to examine the effects of heat stress on same-sex sexual behaviour.

We exposed males to either a benign constant temperature (20°C) or a simulated 3-day heatwave (26°C) and examined the frequency of same-sex sexual behaviour, as well as associated changes in their cuticular profiles. We predicted that (*i*) same-sex sexual behaviour would increase under thermal stress, (*ii*) there would be a divergence in cuticular profiles between thermal treatments, and (*iii*) variation in cuticular profiles under thermal stress would be related to the frequency of same-sex sexual behaviour. More specifically, we expected that heat stress would shift cuticular profiles toward more saturated, longer-chain compounds and that this disruption would be associated with an increased frequency of same-sex mounting due to misrecognition. Such evidence would suggest that mate recognition systems are highly vulnerable to rising temperatures, potentially reducing mating efficiency and reproductive success with implications for population viability.

## Methods

### Animal husbandry

Our laboratory population of beetles originated from wild-caught individuals collected at Tentsmuir National Nature Reserve, UK (56°25′44.7″N 2°52′11.2″W) in June and July 2025. Fourth-generation beetles were reared under controlled laboratory conditions at 20°C in an incubator (LMS Series 1A Model 280 NP) under constant darkness throughout larval and pupal development. Upon eclosion, virgin males were transferred to individual transparent plastic containers (12×8×2 cm) half-filled with moist soil and maintained under the same temperature and light conditions until reaching sexual maturity, approximately 10 days post-eclosion. Beetles were fed raw organic beef twice a week from the time of eclosion and throughout the 3-day temperature treatment.

### Experimental design

Males of a standardised age (10-12 days post-eclosion) were randomly assigned to one of two thermal treatments: (*i*) a control treatment where they continued to be exposed to a constant temperature of 20°C or (*ii*) a simulated heatwave treatment where males were exposed to 26°C for 72 hr[sensu 62]. These conditions reflect ecologically relevant thermal regimes that beetles experience in central Scotland (Met Office 2026).

### Behavioural observations and measurements

At the end of the 72-hour simulated heatwave, we paired up unrelated males from the same thermal treatment (n_control_=32, n_heatwave_=29). One male of each pair was randomly selected and marked on the pronotum with Tipp-Ex, while the other male was lightly poked with a pencil to account for any effect of the marking procedure. Pairs were given one hour to recover from the marking procedure, and subsequently placed in an arena, a 17x12x6 cm transparent plastic container, for behavioural observations. Each arena was filled with approximately 1cm of moist soil, allowing beetles to dig while keeping interactions visible to the observer. Observations were conducted under red light at the same temperature as the thermal treatments (20°C or 26°C).

Behaviours were recorded using focal sampling. Each pair was given one minute to recover after being placed in the arena and then observed for 15 minutes. We recorded three categories of behaviours: “same-sex sexual behaviour” defined as mounting, “aggressive behaviour”, which included grappling and biting, and “overall activity” which combined aggressive and sexual behaviours, plus walking, flying, digging, and self-grooming. For sexual and aggressive events, we measured the frequency and duration of behaviours. For pairs exhibiting same-sex sexual behaviour, we also recorded the latency to its first occurrence. Behaviours involving an interaction were attributed only to the beetle performing the action, not to the recipient (e.g., during mounting, only the mounting beetle was recorded as exhibiting same-sex sexual behaviour). Reciprocal mounting was defined as both males in a pair mounting at least once. After each behavioural trial, both males were immediately euthanised by freezing at -20°C.

### Solvent wash and GC-MS analysis

Prior to extraction, beetles were thawed for about 15 min at room temperature, and Tipp-Ex was scraped from their pronotum to prevent contamination of the cuticular lipid extract. Under a fume hood, each male was individually immersed in 3 mL of n-hexane in a 5 ml glass vial for 15 minutes to dissolve their cuticular compounds. The beetle was then removed, and the solvent extract was allowed to evaporate overnight. Dried samples were stored at -20°C until GC-MS analyses. Following cuticular lipid extraction, we measured pronotum width using digital calipers as a proxy for body size (Pilakouta et al. 2018).

Prior to GC-MS analysis, samples were resuspended in 200 µL of cyclohexane, to which 10 µL of *n*-eicosane (200 ng/µL in cyclohexane) was added as an internal standard. Samples were analysed on a GC-MS (Shimadzu GC2030 gas-chromatograph connected to a Shimadzu QP2020NX mass-spectrometer; Shimadzu, Duisburg, Germany). The GC contained a non-polar capillary column (Rtx-5MS, length = 30 m, inner diameter = 0.25 mm, film thickness = 0.25 µm, Restek, Bellefonte, PA, USA) and used helium as carrier gas with a constant linear velocity of 50 cm/s. The oven temperature started at 40 °C for 2 min and was then raised at a rate of 5 °C/min to 300°C which was held for 6 min. The mass spectrometer was operated in electron ionization (EI) mode at 70 eV and scanned the mass range m/z 35-600. We identified n-alkanes through a comparison of their mass spectra and retention time with a reference mixture of alkanes (Sigma-Aldrich, St. Louis, USA). Other cuticular lipids were identified by interpretation of the MS spectrum and comparison of the retention indices with previously identified compounds from *N. vespilloides*[59] and with the NIST database.

### Statistical analysis

All statistical analyses were performed using R version 4.5.3 (R Core Team, 2026). Generalised linear models were fitted using the *glmmTMB*[65] and *MASS* packages[66], and linear and generalised linear mixed models using the *lme4*[67] and *lmerTest* packages[68]. Multivariate analyses of cuticular hydrocarbon profiles (PERMANOVA) were performed using the *vegan* package[69]. Model diagnostics were performed using the *DHARMa* package to assess dispersion, residual structure, and conformity to distributional assumptions[70]. For data manipulation and figures, we used the *dplyr*[71], *ggplot2*[72], *patchwork*[73], *cowplot*[74], *ggrepel*[75], *tydiverse*[76] and *writexl*[77] packages.

#### Effects of heatwave exposure on male behaviour

Behaviours were summed for both males in each pair to produce pair-level measures of mounting frequency and duration, aggressive event frequency and duration, and overall activity. A total of 61 pairs were analysed with 32 in the control treatment and 29 in the heatwave treatment. Reciprocity, duration and latency to mounting were conditional upon mounting having occurred and were therefore restricted to the 48 pairs in which at least one mount was recorded. Likewise, aggression duration was restricted to the 22 pairs in which aggression occurred.

Pair activity constrains the opportunity for social interactions to occur and was therefore included as a covariate in all models, after confirming that it was independent of thermal treatment with a Tweedie GLM with a log link (z = 0.24, p = 0.81). Activity was log-transformed with a unit offset because of two inactive pairs. Then, the contribution of activity was tested by likelihood-ratio test for each response.

Mounting, reciprocal mounting, and aggression occurrence were analysed using binomial GLMs. Discrete variables were analysed using generalised linear models with a Poisson error distribution (mounting frequency) or a negative binomial distribution (aggression frequency). Continuous variables were analysed using a Gamma error distribution with a log link (mount duration) or a Gaussian error distribution (aggression duration, log-transformed). Latency to first mount was compared between treatments using a Wilcoxon rank-sum test. For each pair in which only one male mounted (n = 29), the difference in body size between the mounting and the non-mounting male was tested using a paired t-test.

#### Effects of heatwave exposure on cuticular lipids

We retained all peaks which represented more than 0.5% of the total peak area in at least one individual and removed non-hydrocarbon substances except aldehydes, resulting in a total of 58 peaks (Figure S1). Family and pair were included as crossed random effects in all mixed models (except in the duration and chain-length models where pair explained no variance and was therefore excluded), and family constrained permutations in the PERMANOVA.

Whole-profile differences between treatments were tested with a permutational multivariate analysis of variance (PERMANOVA; adonis2, vegan) on Bray-Curtis dissimilarities of relative amounts (each compound expressed as a percentage of the total cuticular lipid peak area per individual). For all other analyses, each retained peak area was divided by the internal standard (eicosane) and log-transformed to ensure normality. To characterise the main axes of variation in cuticular lipid amounts, a principal component analysis was run on the correlation matrix, retaining the ten components with eigenvalues >1[78]. For each retained component, we tested for a treatment effect using linear mixed models, correcting p-values for multiple testing (Benjamini-Hochberg). As studies in other insects suggest that relative cuticular lipid composition can be important for recognition[44,79], we repeated the principal component analysis with each compound expressed as a percentage of the total cuticular lipid peak area per individual, and tested each retained component for a treatment effect as above. This gave similar results, so we report the internal-standard analysis in the main text.

Compounds were grouped into classes (n-alkanes, methyl-branched alkanes, alkenes and aldehydes) and amounts of each class were compared between treatments using linear mixed models. To test whether heat shifted the profile toward longer-chained compounds, we compared the abundance-weighted mean chain length of n-alkanes between treatments with a linear mixed model. To test the effect of heat stress on specific compounds previously implicated in sex distinction and copulation in *N. vespilloides*, we analysed the male-typical alkenes 7-C23ene and 7-C25ene[59]. We compared the amounts of each compound between treatments using a linear mixed model with treatment as a fixed effect and family as random effect.

Lastly, to test whether cuticular profiles predicted same-sex mounting, we related principal component scores that were affected by thermal treatment to the frequency of being mounted with a Poisson GLMM. We also tested whether the two lipid classes that changed under heat stress, as well as chain length, predicted mounting frequency using Poisson GLMMs.

## Results

### Effects of heatwave exposure on male behaviour

*Same-sex sexual behaviour*―Pairs in the heatwave treatment performed more mounts than pairs in the control treatment (*β* = 0.45, *p* = 0.028; Figure 1A). More active pairs also mounted more often (*β* = 0.84, *p* < 0.001). Mounting occurrence was not affected by thermal treatment (*β* = 0.66, *p* = 0.47) but was strongly predicted by overall activity with more active pairs being more likely to mount (*β* = 2.40, *p* < 0.001). Among the 48 pairs that mounted, the mean duration of a mount did not differ significantly between thermal treatments (*β* = -0.34, *p* = 0.10). More active pairs performed longer mounts (*β* = 0.80, *p* = 0.004). Reciprocal mounting, in which both males of a pair mounted at least once, was more frequent in the heatwave treatment (*β* = 2.18, *p* = 0.002) (Figure 1B). Overall activity did not predict reciprocal mounting (*β* = -0.93, *p* = 0.28), and latency to first mount did not differ between thermal treatments (*W* = 340, *p* = 0.29). For pairs in which only one male mounted, the mounting male tended to be smaller than the non-mounting male (*t* = -3.20, *p* = 0.003).

**Figure 1:**
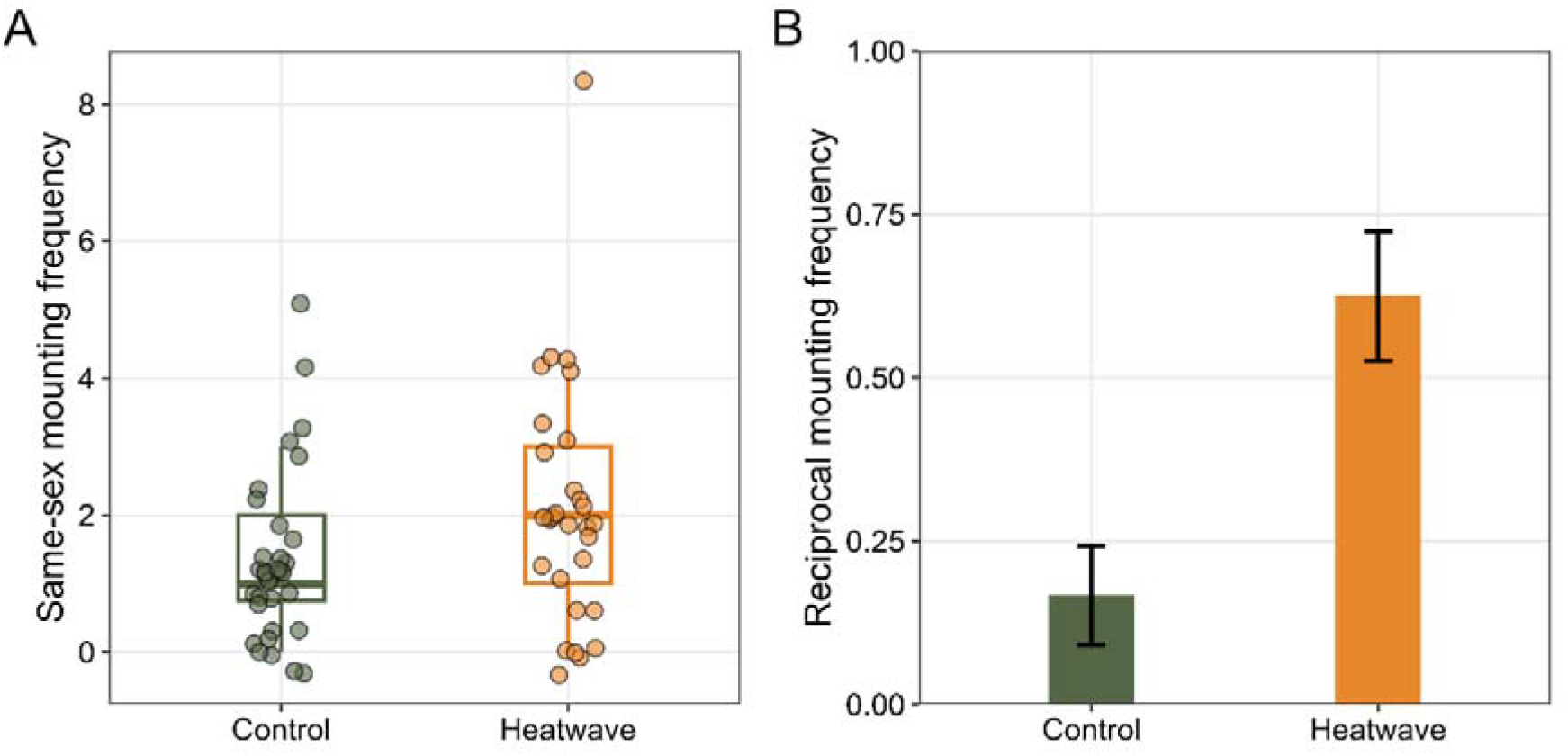
Same-sex mounting frequency in the control and heatwave treatments. (A) Number of same-sex mounts per pair (n_control_ = 32, n_heatwave_ = 29). The lower and upper hinges of the boxes correspond to the first and third quartiles. The whiskers extend from the hinge to the smallest and largest values no further than 1.5 × the interquartile range from the hinge. Points show individual pairs, jittered horizontally to reduce overlap. (B) Proportion of pairs (± SE) showing reciprocal mounting, in which both males mounted at least once (n_control_ = 32, n_heatwave_ = 29).

*Aggression*―Thermal treatment did not affect any measure of aggression. Aggressive interactions occurred in 9 of 32 control pairs and 13 of 29 heatwave pairs, but this difference was not significant (*β* = 0.75, *p* = 0.19). Similarly, neither the number of aggressive interactions (*β* = 0.56, *p* = 0.17) nor their mean duration (*β* = - 0.34, *p* = 0.34) differed between treatments. Overall activity predicted both the occurrence (*β* = 1.19, *p* = 0.046) and the number (*β* = 1.82, *p* = 0.003) of aggressive interactions, but not their mean duration (*β* = 0.15, *p* = 0.68).

### Effects of heatwave exposure on cuticular lipids

The whole cuticular profile differed significantly between control and heatwave beetles (PERMANOVA: R^2^ = 0.13, F = 17.3, *p* = 0.003). Cuticular compounds identified and their relative contribution in heat-stress and control beetles are listed in Table S1. The profile shifted toward saturated and longer-chained compounds. Of the ten PCs retained, only PC3 and PC1 differed significantly between treatments after Benjamini-Hochberg correction (Figure 2a). PC3 contrasted saturated compounds (negative loadings) with unsaturated compounds (positive loadings) (Figure 2b). Heatwave beetles scored lower on this axis, indicating a shift toward more saturated profiles (*β* = -2.76, *p* < 0.0001). PC1 loaded negatively and near-uniformly across almost all compounds. Heatwave beetles scored higher on this axis, indicating lower overall CHC amounts (*β* = 2.75, *p* = 0.030). Expressing compounds as relative composition instead gave the same shift toward more saturated profiles (*β* = -3.13, *p* < 0.0001; Figure S2).

**Figure 2:**
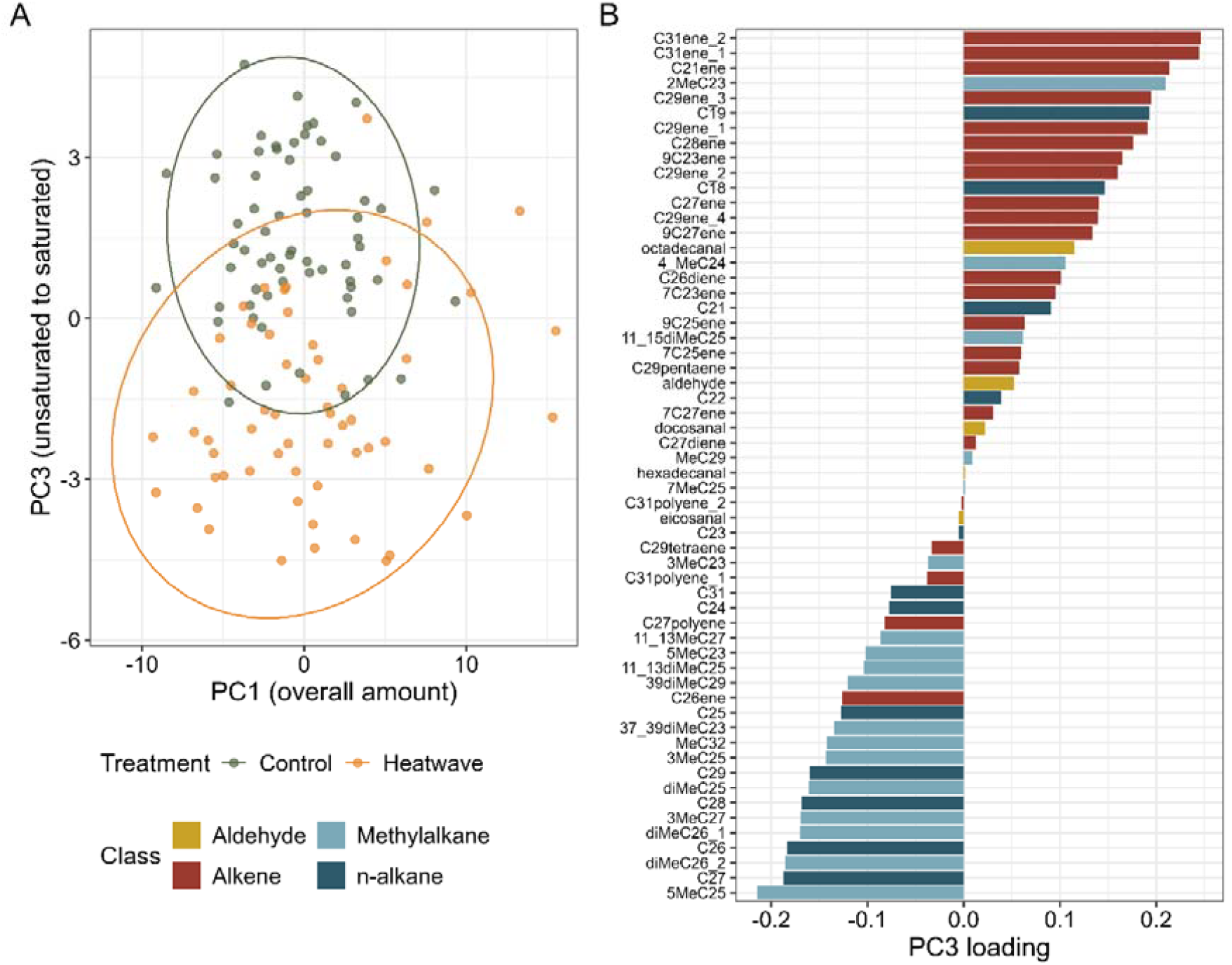
Effect of heatwave exposure on the cuticular profile of male *Nicrophorus vespilloides*. (A) Principal component analysis of internal standard-normalised, log_10_-transformed compound amounts, showing individual beetles on PC1 (overall cuticular lipids abundance; 38.1% of variance explained) and PC3 (saturated-unsaturated axis; 8.9% of variance explained). Ellipses show 95% confidence per treatment. (B) Loadings of each of the 58 compounds on PC3 (n_control_ = 64, n_heatwave_ = 58).

At the compound-class level, alkenes (*β* = -2.6, p < 0.0001) and aldehydes (*β* = - 0.049, *p* = 0.005) were substantially reduced under heat. In contrast, n-alkanes (*β* = - 1.03, *p* = 0.090) and methyl-branched alkanes (*β* = 0.10, *p* = 0.49) did not differ significantly between treatments (Figure 3). In line with our prediction, the abundance-weighted mean chain length of n-alkanes increased significantly under heat (*β* = 0.16, p = 0.004). Among the compounds previously identified as sexually dimorphic, the male-typical alkene 7-C25ene was significantly lower in heat-stressed males (*β* = -0.11, *p* = 0.025), whereas 7-C23ene did not differ between treatments (*β* = -0.0066, *p* = 0.16).

**Figure 3:**
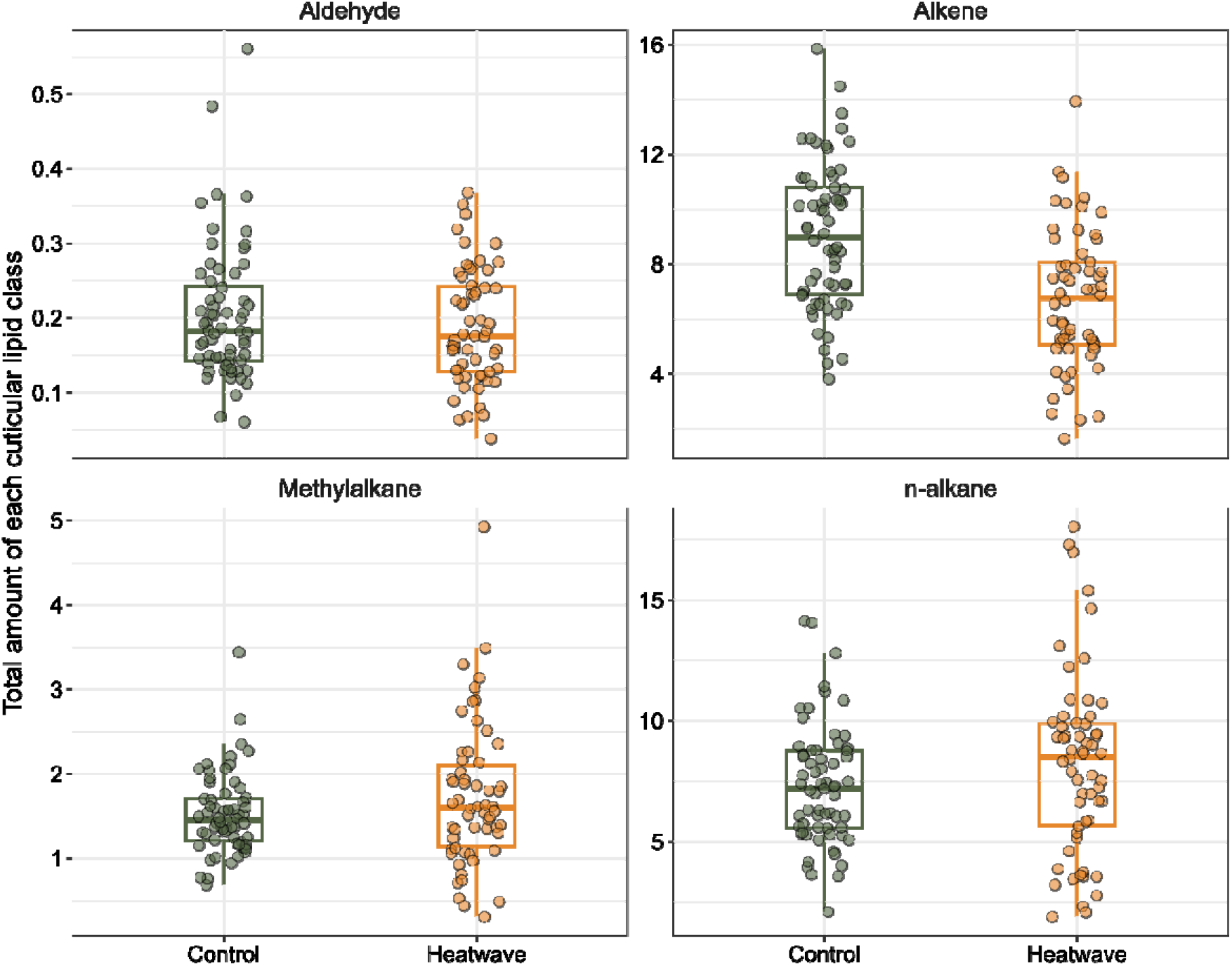
Summed amounts of each cuticular lipid class in control and heatwave male *Nicrophorus vespilloides*, expressed relative to the internal standard. The lower and upper hinges of the boxes correspond to the first and third quartiles. The whiskers extend from the hinge to the smallest and largest values no further than 1.5 × the interquartile range from the hinge. Points show individual beetles, jittered horizontally to reduce overlap (n_control_ = 64, n_heatwave_ = 58).

The shift in cuticular compounds under heat stress did not seem to predict same-sex mounting. Neither of the treatment-affected components predicted the frequency of mounting received (PC1: *β* = 0.012, *p* = 0.63; PC3: *β* = -0.043, *p* = 0.43). Likewise, the amounts of the two compound classes that changed under heat stress―alkenes (*β* = -0.059, *p* = 0.26) and aldehydes (*β* = 0.93, *p* = 0.58)―did not predict mounting frequency. In contrast, n-alkanes chain length did predict the frequency of mounting received (*β* = 0.35, *p* = 0.004) (Figure S3).

## Discussion

In this study, we show that in males of the burying beetle *Nicrophorus vespilloides*, heatwave exposure increased same-sex sexual behaviour and shifted cuticular profiles toward more saturated and longer-chained compounds. We also found some limited evidence that cuticular profiles were related to the occurrence of same-sex mounting. By linking temperature, chemical signalling, and same-sex sexual behaviour, our study offers a new mechanistic perspective on how warming may disrupt sexual communication.

Consistent with our hypothesis, heat stress increased the frequency of same-sex sexual behaviour, as well as the occurrence of reciprocal mounting. This increase in same-sex mounting was not accompanied by more aggressive interactions, which might have been expected if altered information content heightened conflict[81]. On the other hand, if heat stress impairs sex recognition such that males are no longer being identified as male, they would be perceived as less of a threat[82], which is consistent with both the lack of increased aggression and the higher mounting rate we observed. This aligns with prior work identifying mistaken identity as the main explanation for same-sex sexual behaviour in arthropods[83,84]; but see[85].

The observed increase in same-sex sexual behaviour could arise downstream of the cuticular signal produced by the sender, through mechanisms other than misrecognition. At the level of the receiver’s cognitive processing, heat stress can impair learning[86] and memory[87] in insects, and a reduced ability to retain or process social information could increase indiscriminate mounting independently of recognition. Alternatively, as future reproductive prospects decline under thermal stress, individuals may increase investment in current reproduction (terminal investment), as previously observed in this species[63,64]. By making the rejection of a potential mate more costly, this would lower the acceptance threshold and lead to indiscriminate mounting. Interestingly, the mounting male was usually smaller than the male being mounted, which could also reflect a mating filter, where smaller males face greater costs of rejecting a mating opportunity and therefore mount indiscriminately[88,89]. Alternatively, although there is no sexual size dimorphism in our species[90], females are typically larger in elytra length[91], so larger males might be more often mistaken for females, consistent with a misrecognition mechanism.

Temperature-driven behavioural changes were also accompanied by a shift in cuticular compounds, consistent with a functional trade-off. Saturated compounds, especially long-chained, linear ones have higher melting points[92] and should therefore be favoured at high temperatures because they increase desiccation resistance[39]. Our results are in line with this prediction as heat-stressed beetles showed a shift toward more saturated profiles with a reduction in alkenes and an increase in n-alkane chain length. Although specific lipid classes were not related to the frequency of same-sex sexual behaviour, n-alkane chain length was associated with mounting frequency: individuals with longer-chained profiles were mounted more frequently than those with shorter chains. This supports our trade-off hypothesis, since heat stress also shifted profiles toward longer chains.

A closer look at individual compounds revealed contrasting patterns. There is known sexual dimorphism in *N. vespilloides* cuticular profiles: Steiger et al.[59] found that males and females differed mostly in their proportions of the alkenes 7-C23ene and 7-C25ene, which are higher in males. We found that heat-stressed males had less 7-C25ene only, consequently shifting them toward a more female-like cuticular profile, which would be consistent with their being mounted more often. Ambiguous or feminised chemical cues have previously been shown to be subjected to heightened levels of same-sex sexual behaviour (e.g.[93,94]). However, such single compound comparison should be treated cautiously, as sex recognition and courtship are thought to be triggered by the composition of the whole cuticular profile rather than by individual compounds[59,95]. In addition, same-sex mounting under heat stress could also arise from changes in peripheral chemosensory sensitivity or central processing, independent of any shift in the cuticular signal itself[23,29].

At a conceptual level, our work contributes to the long-standing question of how same-sex sexual behaviour arises mechanistically. We show that heat stress simultaneously increases same-sex mounting and alters cuticular profiles, which is consistent with same-sex sexual behaviour arising as a context-dependent outcome, whether through imperfect information or a shift in the acceptance threshold. However, we only found limited evidence that the specific cuticular lipid changes predicted mounting, so the causal link between altered signalling and behaviour remains to be established. Unravelling the mechanistic underpinnings of same-sex sexual behaviour would help move the field beyond purely adaptive-versus-non-adaptive debates toward integrated mechanistic explanations.

Given the conserved role of CHCs in arthropod social communication and their sensitivity to temperature[41,96], we would expect this thermal disruption of chemically mediated recognition and the observed increase in same-sex sexual behaviour to be a common response, at least in arthropods. In addition, temperature has been shown to affect sexual chemical signalling in other taxa, particularly reptiles[97–99]. The effects of temperature on same-sex sexual behaviour may therefore be taxonomically widespread beyond arthropods. We recommend that future work tests for these effects in other species to determine the generality of this pattern.

In summary, our study provides the first evidence for a causal effect of temperature on the rate of same-sex sexual behaviour and offers novel insights into the detrimental effects of warming on animal reproduction due to a disruption in communication. This work thus provides evidence for a proximate, mechanistic explanation (i.e., signal disruption) for same-sex sexual behaviour. Given the conserved role of CHCs and other types of pheromones in social communication and the fact that they are highly sensitive to temperature, we suggest that this pattern may be generalisable across and beyond arthropods. Our findings therefore raise broader questions about how animals that rely on chemical signalling may cope in a warming world.

## Acknowledgements

NP & SM conceived the study, NP, SM, NWB and SS designed the experiment; SM performed the behavioural experiment; JS and SM performed the chemical analyses; SM wrote the first draft of the manuscript; all authors reviewed and edited the manuscript.

We would like to thank Rebekah Douglas, Jakob Wiil, Izzy Grieve, and Sam Wraight for their help with collecting wild-caught beetles and with animal husbandry for the lab population. We also thank Jakob Wiil and Izzy Grieve for their helpful feedback on an earlier version of this manuscript.

## Supplementary figure legends

**Figure S1:** Representative gas chromatogram of the cuticular profile of a male *Nicrophorus vespilloides*. Labelled peaks indicate the cuticular compounds identified. C20 is the internal standard.

**Table S1:** Mean relative contribution (%) (± SD) of each cuticular compound in heat-stressed and control beetles. Compounds are listed in order of increasing retention index.

**Figure S2:** Effect of heatwave exposure on the cuticular profile of male *Nicrophorus vespilloides*. Loadings of each of the 58 compounds on PC2 from the relative-composition PCA (n_control_ = 64, n_heatwave_ = 58).

**Figure S3:** Abundance-weighted mean chain length of n-alkanes in control and heatwave-exposed male *Nicrophorus vespilloides*. The lower and upper hinges of the boxes correspond to the first and third quartiles. The whiskers extend from the hinge to the smallest and largest values no further than 1.5 × the interquartile range from the hinge. Points show individual beetles, jittered horizontally to reduce overlap (ncontrol = 64, nheatwave = 58).

