## Supplementary figures and images for "Mixed signals: heat stress alters chemical profiles and increases same-sex sexual behaviour in an insect"

### Figure S1

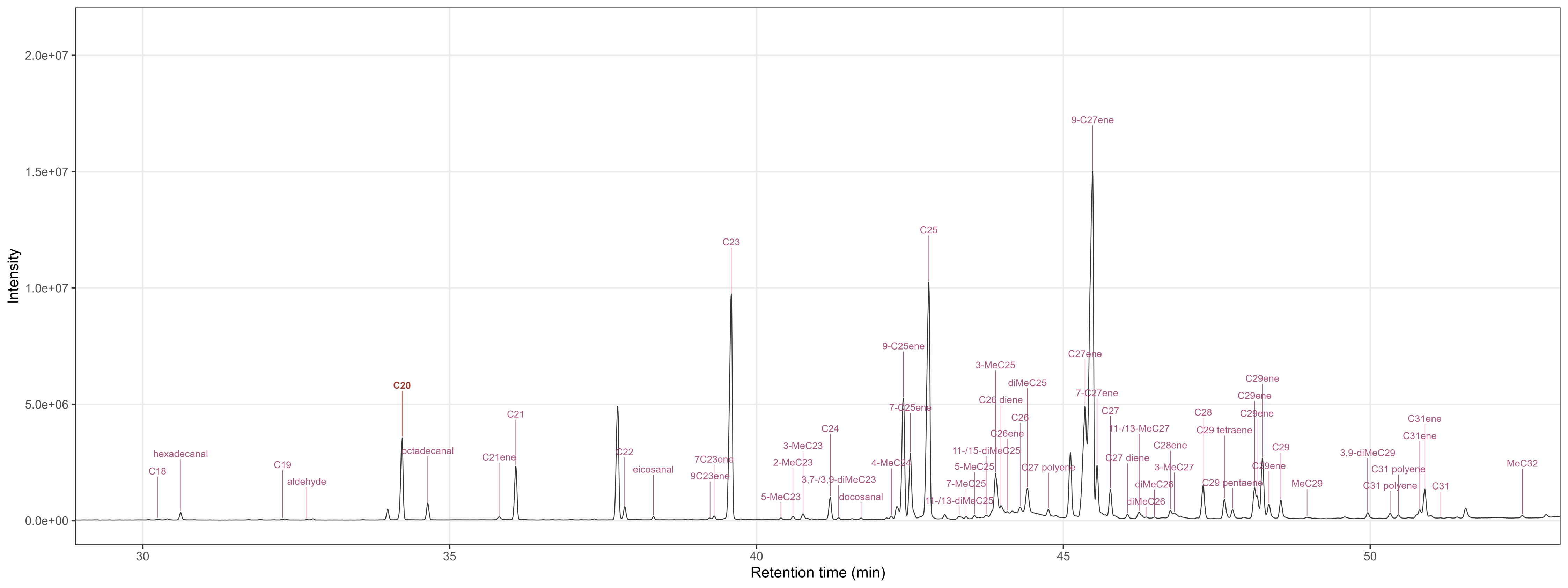

### Figure S2

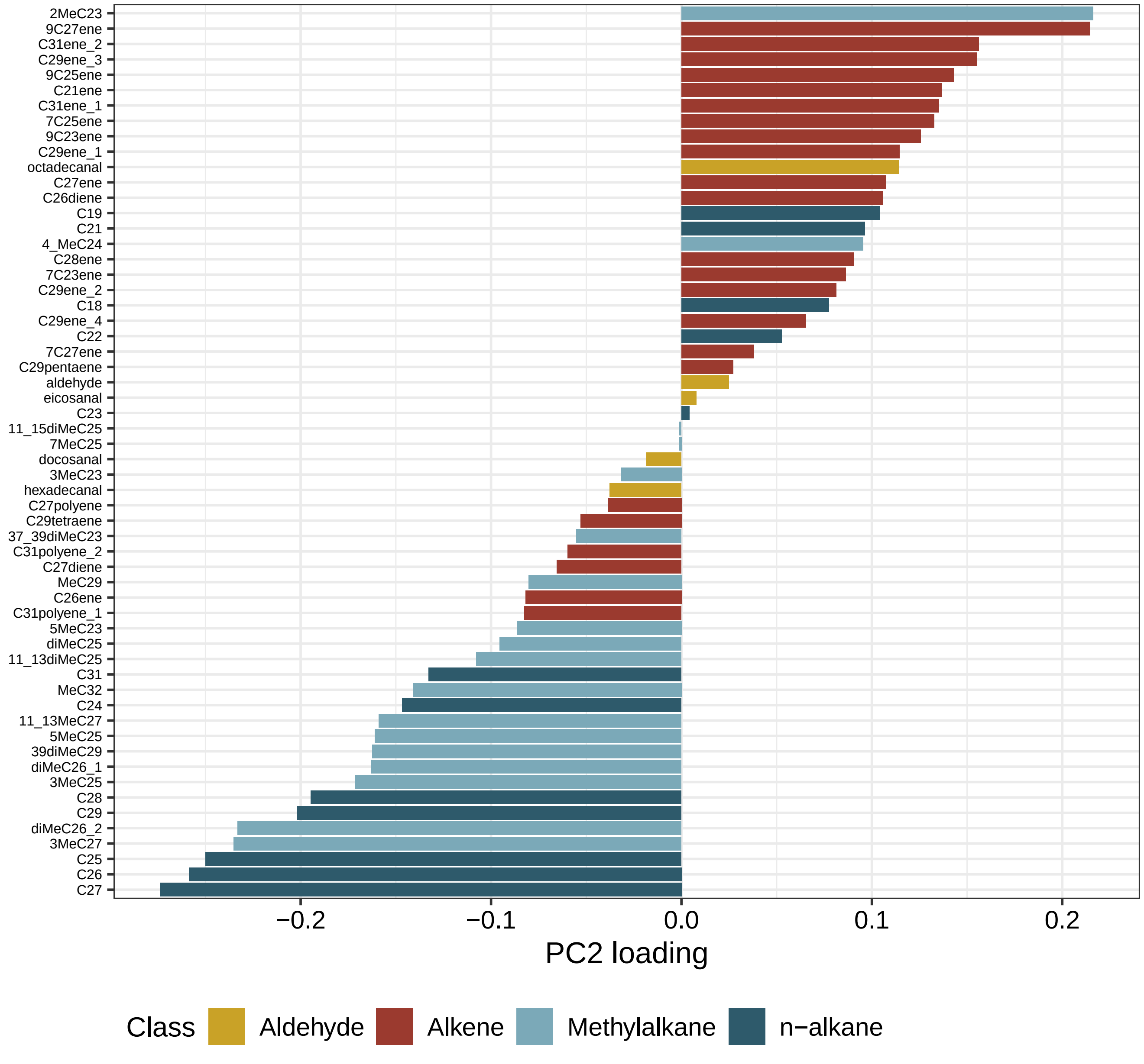

### Figure S3

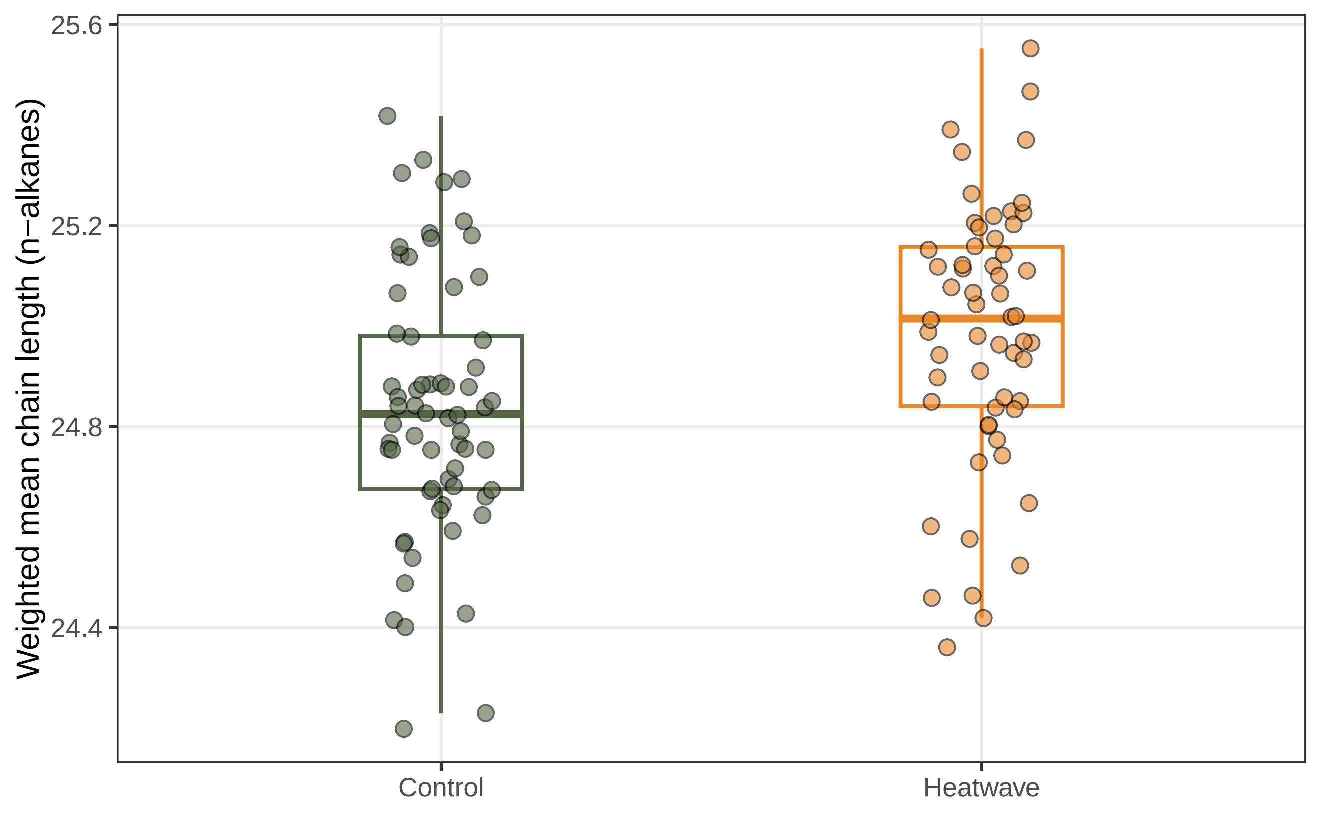
