## Supplementary material for "Mixed signals: heat stress alters chemical profiles and increases same-sex sexual behaviour in an insect": Table S1

| compound | mean control | SD control | mean heat | SD heat |
|---|---|---|---|---|
| C18 | 0,037958865 | 0,032309886 | 0,049436826 | 0,067978795 |
| hexadecanal | 0,342889965 | 0,200458492 | 0,372441308 | 0,243900946 |
| C19 | 0,019303204 | 0,009907956 | 0,020011157 | 0,016882657 |
| aldehyde | 0,015867342 | 0,006105763 | 0,015832854 | 0,006833613 |
| octadecanal | 0,474477308 | 0,167389138 | 0,486339053 | 0,237230206 |
| C21ene | 0,158413854 | 0,107565523 | 0,087197019 | 0,069738162 |
| C21 | 0,815849371 | 0,62946495 | 0,700988927 | 0,669202126 |
| C22 | 0,381674461 | 0,197369788 | 0,403127879 | 0,270646361 |
| eicosanal | 0,131464378 | 0,06287517 | 0,150710839 | 0,064170478 |
| 9C23ene | 0,07592043 | 0,082573782 | 0,060495964 | 0,077623645 |
| 7C23ene | 0,153803932 | 0,099923717 | 0,140382154 | 0,122635971 |
| C23 | 9,926512565 | 3,082090101 | 10,36893129 | 3,12104064 |
| 5MeC23 | 0,077571574 | 0,053039592 | 0,108764769 | 0,06949316 |
| 2MeC23 | 0,231566811 | 0,073790159 | 0,15282253 | 0,070593021 |
| 3MeC23 | 0,273718059 | 0,102766157 | 0,312507182 | 0,149864957 |
| C24 | 1,722758383 | 0,39542138 | 1,912624618 | 0,475461109 |
| 37_39diMeC23 | 0,04005569 | 0,025226092 | 0,06936281 | 0,075077875 |
| docosanal | 0,153582017 | 0,083493106 | 0,159314268 | 0,068033286 |
| 4_MeC24 | 0,239415008 | 0,076964686 | 0,172001829 | 0,069203128 |
| 9C25ene | 2,064811971 | 1,874074104 | 1,688053241 | 1,740775729 |
| 7C25ene | 2,474794494 | 1,260875349 | 2,096706608 | 1,322143475 |
| C25 | 19,88148933 | 3,627329811 | 24,11714856 | 4,511833769 |
| 11_13diMeC25 | 0,119275661 | 0,04506478 | 0,18073766 | 0,103264856 |
| 7MeC25 | 0,096901099 | 0,031815584 | 0,106684774 | 0,047645407 |
| 5MeC25 | 0,171641719 | 0,07447205 | 0,281747833 | 0,126549827 |
| 11_15diMeC25 | 0,33206729 | 0,129131933 | 0,313605376 | 0,202413318 |
| 3MeC25 | 3,121034706 | 0,8961389 | 3,599086041 | 0,995994422 |
| C26diene | 0,165531783 | 0,058712981 | 0,142972977 | 0,074283728 |
| C26ene | 0,037072558 | 0,044105033 | 0,055855754 | 0,043704202 |
| C26 | 0,658168391 | 0,209950108 | 0,987549897 | 0,418072518 |
| diMeC25 | 1,118086761 | 0,378883618 | 1,562880662 | 0,899131415 |
| C27polyene | 0,087668404 | 0,078807478 | 0,122809122 | 0,087464392 |
| C27ene | 4,299212052 | 1,309117071 | 3,499815336 | 1,11718791 |
| 9C27ene | 21,07425265 | 3,854563152 | 18,49228914 | 3,419877147 |
| 7C27ene | 2,701140278 | 1,114515953 | 2,524917104 | 0,708307806 |
| C27 | 2,806446368 | 0,872083143 | 4,395879459 | 1,818097041 |
| C27diene | 0,430800073 | 0,282571981 | 0,52371127 | 0,31291812 |
| 11_13MeC27 | 0,537874096 | 0,161025866 | 0,696879155 | 0,231417679 |
| diMeC26_1 | 0,052017752 | 0,025014226 | 0,084581272 | 0,068204163 |
| diMeC26_2 | 0,141937856 | 0,058702208 | 0,258487912 | 0,115330445 |
| C28ene | 0,347203453 | 0,139230871 | 0,223726954 | 0,101802931 |
| 3MeC27 | 0,43767312 | 0,141014861 | 0,590071824 | 0,189077533 |
| C28 | 2,789278905 | 0,890587594 | 3,305310994 | 1,174868958 |
| C29tetraene | 0,841926957 | 0,330789148 | 1,107138803 | 0,591233945 |
| C29pentaene | 0,487603708 | 1,032190192 | 0,91478851 | 2,637720805 |
| C29ene_1 | 1,337696192 | 0,564913032 | 0,843147772 | 0,390787141 |
| C29ene_2 | 1,014211186 | 0,368606607 | 0,800711287 | 0,420378426 |
| C29ene_3 | 4,347217292 | 1,376893009 | 2,964636381 | 1,279624191 |
| C29ene_4 | 2,156265636 | 1,194595072 | 1,415351957 | 0,799942381 |

| | | | | |
|---|---|---|---|---|
| C29 | 1,145986644 | 0,470322822 | 1,586029089 | 0,66723911 |
| MeC29 | 0,217288563 | 0,081797278 | 0,224478348 | 0,112841607 |
| 39diMeC29 | 0,874708733 | 0,361462246 | 0,890055091 | 0,33463947 |
| C31polyene_1 | 0,235985638 | 0,084385374 | 0,290109509 | 0,150718627 |
| C31polyene_2 | 0,145626637 | 0,048233466 | 0,180608932 | 0,093647552 |
| C31ene_1 | 1,066309742 | 0,622716858 | 0,478235159 | 0,392554296 |
| C31ene_2 | 4,384820736 | 2,427569373 | 2,076890115 | 1,945375376 |
| C31 | 0,035371196 | 0,0181486 | 0,050255647 | 0,036043435 |
| MeC32 | 0,489797152 | 0,240323705 | 0,58276121 | 0,230279673 |
